# Revealing Heavy Metal Resistances in the Yanomami Microbiome

**DOI:** 10.1101/2023.05.04.539487

**Authors:** Liliane Costa Conteville, Joseli Oliveira-Ferreira, Ana Carolina P Vicente

## Abstract

**BACKGROUND:** The Amazon hosts invaluable and unique biodiversity as well as mineral resources. As a consequence, there are large illegal and artisanal gold mining areas in indigenous territories. Mercury has been used in gold mining, and some are released into the environment and atmosphere, primarily affecting indigenous as the Yanomami. In addition, other heavy metals have been associated with gold mining and other metal-dispersing activities in the region.

**OBJECTIVE:** Investigation of the Yanomami gut microbiome focusing on metal resistance.

**METHODS:** Metagenomic data from the Yanomami gut microbiome were assembled into contigs, and their putative proteins were matched to a database of metal resistance proteins.

**FINDINGS:** Most identified proteins have the potential to confer resistance to multiple metals (two or more), followed by mercury, copper, zinc, chromium, arsenic, and others. Operons with potential resistance to mercury, arsenic, chromium, nickel, zinc, copper, copper/silver, and cobalt/nickel were identified. Mercury resistance operon was the most abundant, even though a diversity of operons in the Yanomami microbiome was observed to have the potential to confer resistance to various metals

**CONCLUSION:** The Yanomami gut microbiome gene composition shows that these people have been exposed directly or indirectly to mercury and other heavy metals.

**Sponsorships:** This study was partly financed by the Fundação de Amparo à Pesquisa do Estado do Rio de Janeiro (FAPERJ); and PAEF (IOC-023-FIO-18-2-47).

## Introduction

Heavy metals are naturally occurring elements found in the environment ^(1)^. And in biological systems, they play an essential role in carrying out various physiological and biochemical functions ^(2,3)^. However, the distribution and amount of heavy metals have increased significantly worldwide due to various anthropogenic activities such as mining, industrial practices, and the addition of metals to fertilizers and livestock feed ^(4,5)^. Thus, high concentrations of heavy metals have been detected in soils, sediments, and water bodies. The resulting selection pressure often leads to dramatic changes within their microbial populations and ecological functions ^(6–11)^. This process has exceeded the tolerable baseline level of natural heavy metal cycling in many ecosystems and is currently a problem not only for the environment but also for public health ^(2)^.

Exposure of humans and animals to heavy metals can occur through water, air, food, or skin, and bioaccumulation of heavy metals can lead to various toxic effects on different tissues and organs ^(12–14)^. Arsenic, cadmium, and chromium, for example, are considered carcinogenic metals that can impair DNA synthesis and repair ^(15,16)^. To cope with high metal concentrations, microbes have evolved adaptation and resistance mechanisms such as metal resistance (MR) genes ^(17)^. These mechanisms are thought to result from billions of years of selection by geological heavy metals and later by anthropogenic influences ^(2,18)^.

The Amazon hosts an invaluable and unique biodiversity of flora and fauna on Earth and is home to 173 ethnic groups, with indigenous territories occupying a significant portion of the region (27% of the forest area) ^(19)^. Nevertheless, the Brazilian Amazon is home to the third largest area of artisanal mining in indigenous territories ^(20)^, where mercury (Hg) is used to combine with gold in the amalgamation process ^(21)^. Mercury is considered one of the ten most hazardous substances to health by the World Health Organization (WHO) due to its high toxicity ^(22)^. And in the amalgamation process, some mercury is dispersed into rivers, soils, and the atmosphere ^(23,24)^, affecting indigenous areas such as the Yanomami ^(23)^. The Yanomami are the largest indigenous, semi-isolated group in the Amazon that still maintains traditional subsistence practices (hunting, fishing, gathering, and swidden horticulture). Their land encompasses the Venezuela-Brazil border and spans two Brazilian states (Roraima and Amazonas) ^(24)^. Reports indicate that this group has been exposed to mercury from gold mining since 1980 ^(25)^, and it is a current and growing threat ^(26)^. The Yanomami are even more vulnerable because of their dependence on resources in this potentially contaminated environment. Fish in the Amazon, the main source of protein for traditional groups, is also contaminated with mercury ^(27,28)^. And in fact, dietary exposure is the major pathway for the accumulation of metals in the body. In addition to mercury, other heavy metals such as copper (Cu), zinc (Zn), arsenic (As), cadmium (Cd), and lead (Pb) have also been associated with gold mining ^(29)^. However, other metal-dispersing activities may also be occurring in the region. As our group has previously noted, metal contamination could result from the continuous discharge of batteries by the Yanomami on their land for decades ^(30)^.

In addition to the toxic effects caused by metal contamination, the gut microbiota of individuals may reflect their long-term metal exposure ^(31)^. However, studies of this nature are extremely limited, and we found none that focused on isolated or semi-isolated groups. Thus, to gain insight into this issue concerning the Brazilian Yanomami people, we surveyed their gut microbiome, focusing on the heavy metal resistance-associated genes.

## Materials and Methods

### Data Used

For this study, we analyzed shotgun metagenomic data (n = 15) from the gut microbiome of a Brazilian Yanomami group, a semi-isolated hunter-gatherer community from the Brazilian Amazon ^(30)^. The metagenomic data comprises paired-end sequences generated on Illumina sequencer and can be found in the National Center for Biotechnology Information Sequence Read Archive (https://www.ncbi.nlm.nih.gov/sra) under the BioProject PRJNA527208.

### Metagenomic Assemble and Open Reading Frames Prediction

Each metagenome was independently subjected to *de novo* assembly through metaSPAdes v.3.13 ^(32)^ using default parameters and *k-mers* of size (21, 33, 55). This step assembled the short reads of the metagenomes into 2,012,469 contigs (longer sequences). Open Reading Frames (ORFs) were predicted in these contigs with FragGeneScan v.1.31 ^(33)^ using the parameters “-w 1 -t complete”.

### Sequences Classification

The translated ORFs were matched to specific proteins using the Abricate program (https://github.com/tseemann/abricate). This program uses BLAST+ ^(34)^ to compare sequences with reference databases, and in this study, we used the BacMet2 database ^(17)^ to analyze metal resistance proteins. The results from each search were filtered, so only proteins with more than 80% coverage and 80% identity were considered for further analysis.

To relate the identified proteins to the bacterial species of the Yanomami/Brazil microbiome, the contigs that showed similarities to MR proteins were taxonomically classified using the Kraken2 program ^(35)^.

### Metals Determinants Operons

To identify the presence of metal resistance operons, we surveyed the presence of coding genes in the metagenomes. Mercury resistance operon was determined by the *mer* operon, which consists of genes responsible for mercuric reduction (*merA*), periplasmic binding (*merP*), transport (*merTCEG*), and transcriptional regulation (*merRD*) ^(36)^, but also genes associated with organomercury compounds degradation (*merB* and *merG*) ^(37)^. Arsenic resistance operon was determined by the ars operon (inorganic arsenic), which has many variants but, at the minimum, it consists of genes responsible for arsenite efflux transporter ATPase subunit (*arsB*), arsenate reductase (*arsC*) and a metalloregulatory transcriptional repressor (*arsR*) ^(38)^. The copper resistance operon was determined by a system for copper efflux (*cue*) and a system for copper sensing (*cus*) ^(39)^. The cue system consists of the periplasmic multicopper oxidase (*cueO)*, the membrane-embedded ATP-driven copper efflux pumps (*copAB*), and the transcriptional regulator (*cueR)* ^(39,40)^. The cus system has the potential to also confer Silver resistance, and it includes a complex of proton gradient–driven efflux pump (*cusABC*) and regulators (*cusRS*) ^(39)^. The zinc resistance operon was determined by the presence of proteins that control zinc homeostasis by importing and exporting Zn ions via Zn uptake systems ^(41)^. The proteins *znuABC* are ABC transporters that facilitate zinc ions uptake, which can be regulated by *zitB, zur*, or soxS ^(41,42)^. Chromate resistance operon was determined by the chromate ion transport (*chr*) superfamily, which consists of genes responsible for reduction (*nfsA* and *chrR/yieF*), transport (*chrA*), regulation (*chrB* and *chrF*) and superoxide dismutase (*chrC*) ^(43)^. Nickel resistance operon was determined by the *nik* system of ABC transporters, which is involved in specific nickel uptake and is composed of a periplasmic binding protein (*nikA*), integral membrane components (*nikBC*), two ATPases (*nikDE*), and a repressor that avoids nickel overload ^(44,45)^. Cobalt and nickel resistance operon was determined by the *rcn* system, which is composed of a regulator (*rcnR*) that controls the transcription of cobalt and nickel exporters (*rcnAB*) ^(46)^.

## Results

We reanalyzed the metagenomic data from the gut microbiome of Yanomami individuals that inhabit the Brazilian Amazon (Figure 1).

**Figure 1:**
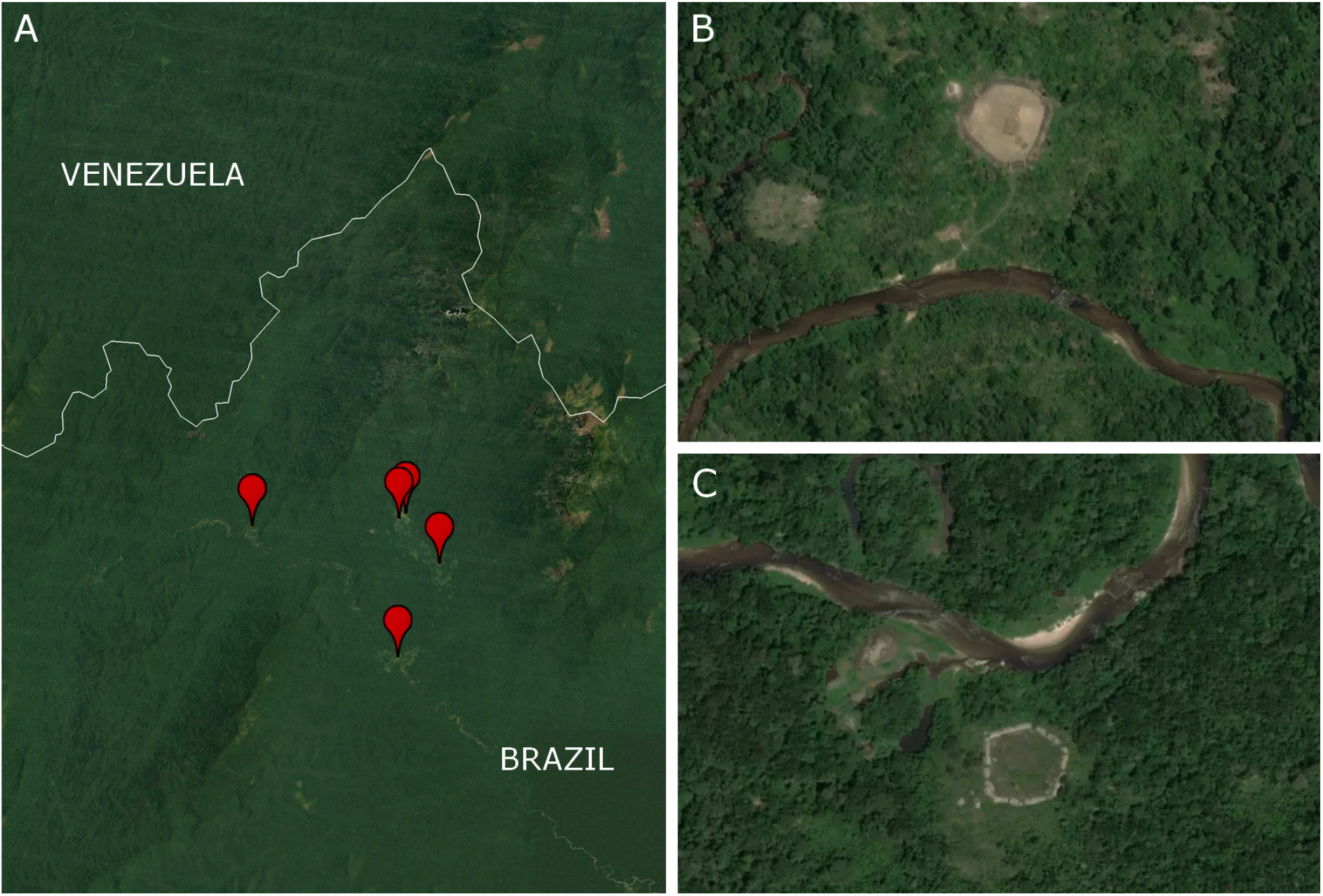
Yanomami villages in the Brazilian Amazon. Satellite images showing (A) part of the large territory explored by the Yanomami and the locations of the villages, and (B-C) two villages near the Marari River. Source: Google Earth and Earthrise Media (https://github.com/earthrise-media/mining-detector).

Putative proteins from the metagenomic contigs were screened for similarity with metal resistance (MR) proteins from the BacMet2 database. In total, 264 translated ORFs showed similarity with 107 MR proteins, and these ORFs are distributed into 174 contigs from 11 of the 15 metagenomes (Figure 2A). Most of these MR proteins have the potential to confer resistance to multiple metals (two or more metals), followed by mercury, copper, and zinc (Figure 2B).

**Figure 2:**
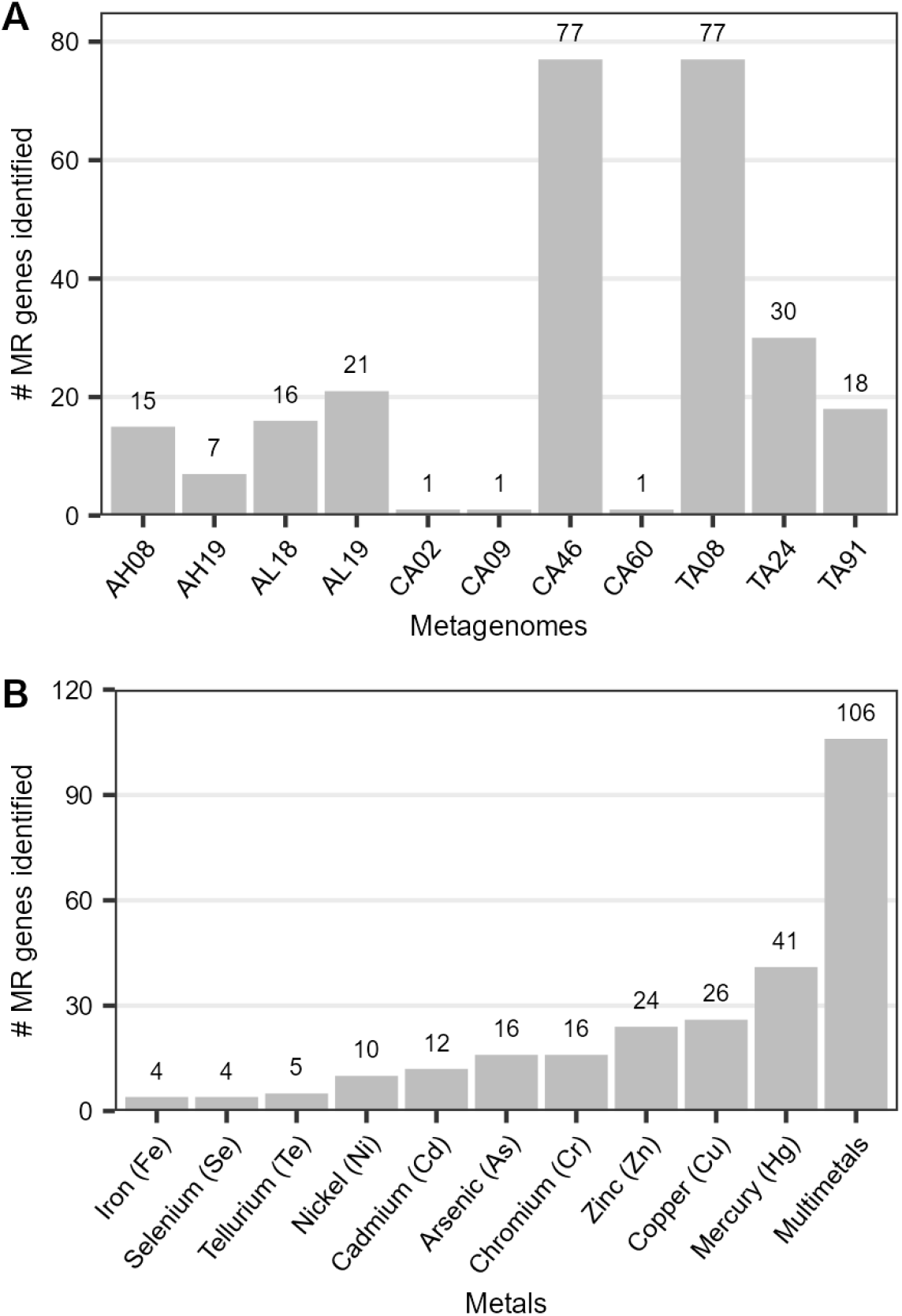
Amount of MR genes identified per (A) metagenome and (B) per metal.

Proteins with more copies in our data are encoded by merT and merP (identified eleven and nine times, respectively), followed by pstB and chrA1 (identified eight times). *merT* was identified in four metagenomes, having up to five copies each, while merP was identified in five metagenomes with up to three copies each. *pstB* was identified in six metagenomes, duplicated in two of them. *chrA1* was duplicated in the four metagenomes in which it was identified.

Contigs that potentially encode MR proteins belong to the phylum *Proteobacteria* (170 contigs) and *Firmicutes* (4 contigs). Of the phylum *Proteobacteria*, the most abundant genus was *Escherichia* (114 contigs with 80 MR genes), followed by *Ralstonia* (18 contigs with 7 MR genes). Of the phylum *Firmicutes*, the genera identified were *Enterococcus* (3 contigs with 4 MR genes) and *Streptococcus* (1 contig with 1 MR gene).

We further investigated the presence of heavy metals and multimetal operons (Figure 3) and identified six operons or systems (*mer, ars, chr, nik, znu*, and *cue*) that comprise genes with resistance potential for mercury, arsenic, chromium, nickel, zinc, and copper, respectively. We also identified two operons (*cus* and *rcn*) associated with multiple metals (copper/silver and cobalt/nickel).

**Figure 3:**
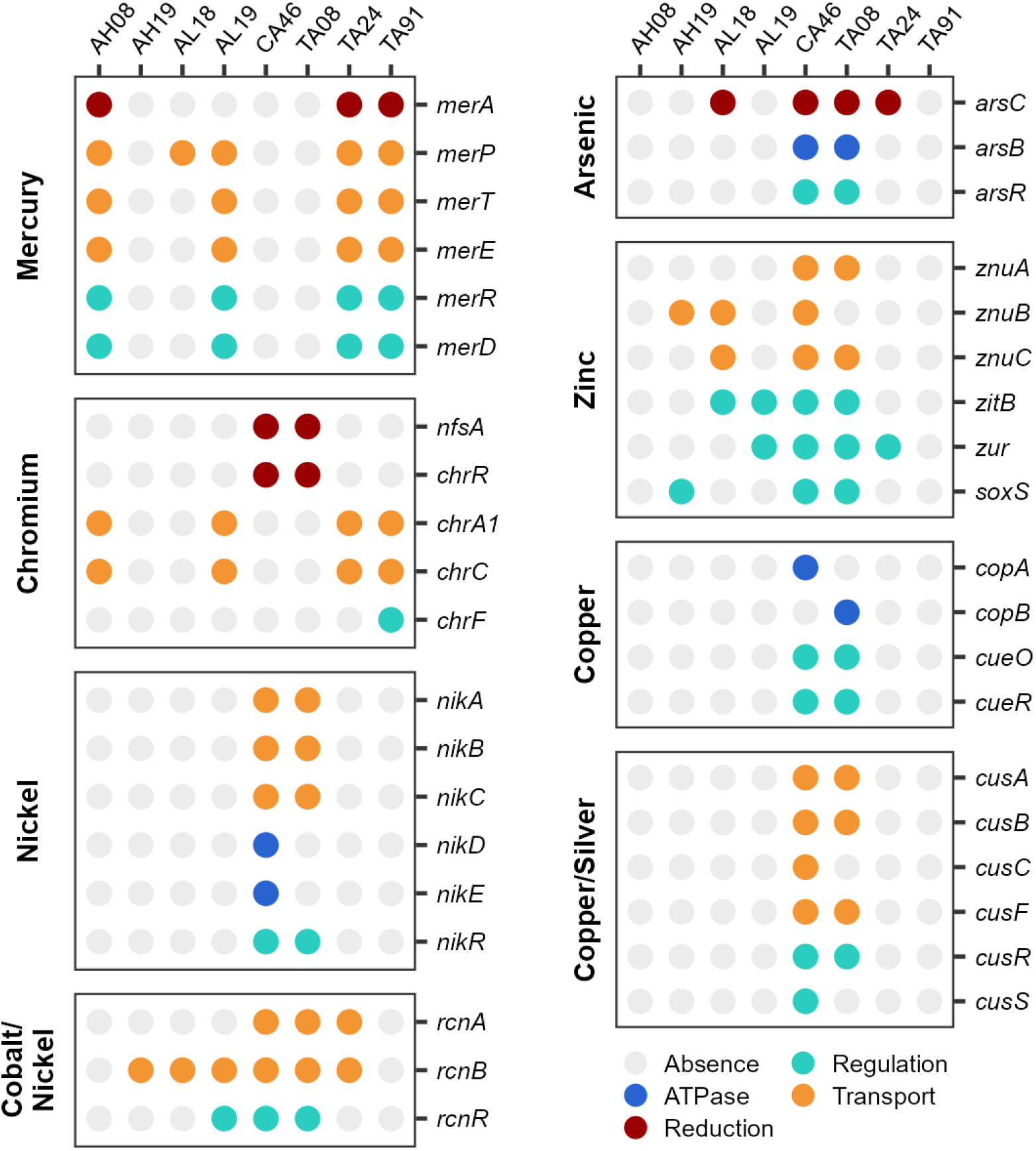
Dot chart representing the genes identified in each metagenome colored by function and grouped by the metal(s) in which they have the potential to confer resistance

The *mer* operon was the most predominant in the metagenomes, being identified in AH08, AL19, TA24, and TA91, which have coding genes for proteins responsible for the main functions necessary for mercury resistance, but the mechanisms for organomercury compound degradation (*merB* and *merG*) were not identified (Figure 3). All contigs harboring genes from the *mer* operon were classified as *Proteobacteria*, mainly *Aeromonas, Burkholderia*, and *Paraburkholderia*. These four metagenomes also harbor genes that encode proteins involved in chromium transport (*chrA1* and *chrC*) in contigs classified as *Ralstonia*. In contrast, the metagenomes CA46 and TA08 harbor genes that encode proteins involved in chromium reduction (nfsA and chrR) in contigs that were classified as *Escherichia, Ralstonia, Paraburkholderia, Kinneretia*, and *Enterobacteriaceae* (Figure 3).

Moreover, the metagenomes CA46 and TA08 stand out for having the highest number of operons detected (nickel, cobalt/nickel, arsenic, zinc, copper, and copper/silver). Most of the contigs (64/70) harboring genes from these operons were classified as *Escherichia coli*. The same contig from metagenome CA46 harbors the complete *nik* and *ars* operons, while the genes from the *rcn* and *cus* operons were identified in different but also single contigs.

## Discussion

In general, the structure, composition, and diversity of gut microbiomes reflect the living conditions and environment of the individuals ^(47)^. Although heavy metals are naturally occurring elements found mainly in soil, heavy metal pollution in mining areas is a serious environmental and human health concern. In our study, we characterized heavy metal resistance genes/proteins associated with the gut microbiome of Yanomami individuals. The Yanomami are a semi-nomadic hunter-gatherer group from the Amazon. Although they explore a rich and highly biodiverse territory, gold mining has been practiced in their land for decades ^(25)^. This practice usually relies on a process that results in the dispersal of mercury into the atmosphere and environment ^(48,49)^. Here, the identification of the *mer* operon and other heavy metal-associated operons in the Yanomami microbiome suggests that they have been simultaneously exposed to mercury and other heavy metals, which may have cumulative effects and cause higher toxicities ^(13)^.

Most identified proteins have been associated with resistance to toxic metals such as mercury, arsenic, and chromium ^(14)^. The genes (*merT* and *merP*) required for the full expression of mercury resistance ^(50)^ had the largest number of copies in the metagenomes, and furthermore, the operon associated with mercury resistance was the predominant one in the metagenomes. The narrow-spectrum mercury resistance merRTPADE operon confers resistance to inorganic mercury, while the broad-spectrum mercury resistance merRTPAGBDE operon confers resistance to inorganic and organic mercury. And both include the central enzyme in the microbial mercury detoxification system: the mercuric reductase (MERA) protein, which catalyzes the reduction of Hg(II) to volatile Hg(0). Organic mercury compounds are more toxic than inorganic compounds because they are readily absorbed from the gastrointestinal tract (95%) and distributed throughout the body ^(14)^. But all forms of mercury are toxic and have been associated with neurological damage, renal dysfunction, gastrointestinal ulceration, and hepatotoxicity in humans ^(14,51)^.

Other abundant genes identified in the Yanomami microbiome encode the proteins CHRA1 and PSTB. The former reduces chromate accumulation in cells, and the latter is involved in phosphate import and arsenic resistance. Bioaccumulation of both metals in the human body can cause dermal, renal, neurological, and gastrointestinal diseases and several cancers. Arsenic, however, is of particular concern because it can endanger health even at very low concentrations and affect the cardiovascular, hepato-biliary, and respiratory systems, in addition to those already mentioned ^(14,51–53)^. In the Amazon forest, chromium concentrations in a pristine area have been reported to be higher than the background concentration, but its origin was presumed to be geogenic ^(54)^. A study from 2005 reported that arsenic concentrations in Amazon rivers were higher in those flowing from the Andes, which is not the case for rivers to which the Yanomami have access ^(55)^. Since then, however, gold mining in the Yanomami region has increased substantially ^(26)^, and elevated arsenic concentrations in sediments, soils, and water in other regions of Brazil have been linked to various anthropogenic activities ^(56)^.

For decades, the Yanomami ecological niche has been under the continuous discharge of batteries that also contributes to releasing metals into their environment ^(30)^. The most commonly used battery types contain significant amounts of heavy metals whose resistance genes were identified in our study, such as cobalt, copper, and nickel ^(57)^. Interestingly, cobalt and copper, as well as zinc, are essential for human life. However, they can also be toxic when present in excess ^(2)^.

As for the bacteria carrying these MR genes, most contigs were classified as *Proteobacteria*. This phylum is widespread and ubiquitous, and due to its adaptability and tolerance, it has been reported to harbor a variety of MR determinants in different environments and contexts ^(2)^. Interestingly, we have previously observed that the gut microbiome of Yanomami individuals differs from that of other semi-isolated human groups in the high abundance of *Proteobacteria* ^(30)^. However, it is not known whether this is the reason why these individuals have more *Proteobacteria* in their gut microbiome. In fact, we have previously linked this high abundance to the high exposure of the Yanomami to solar UVB light ^(58)^.

In summary, identifying genes that encode proteins associated with metal resistance in the feces of Yanomami individuals suggests a high abundance of heavy metals in the ecological niche they explore. Both geological and anthropogenic heavy metals are widespread in the environment and non-degradable, which may constitute long-term selection pressures ^(1)^. Therefore, we cannot trace the source or time period of exposure with this study, but the evidence shows that increases in gold mining have direct impacts on this group, as do increases in deforestation rates ^(20,26)^. Monitoring and remediation of these activities are urgently important not only for the health and survival of indigenous groups but also for the conservation of the Amazon region overall, which implies preserving global biodiversity and regulating the climate and hydrological cycle ^(59)^.

## Acknowledgements

We are grateful to the Yanomami people, and we also thank the health personnel of the Distrito Sanitário Especial Indígena Yanomami for their overall support during fieldwork.

## Author’s contribution

JO-F collected the samples. LC processed the samples and analyzed the data. LC and AV interpreted the data and drafted the manuscript. All authors revised the manuscript, approved the final version to be published, and agreed to be accountable for the work.

